# Heterotopic ossification in intact rat Achilles tendons is characterized by unique mineralized collagen fiber structures

**DOI:** 10.1101/2022.06.28.497706

**Authors:** Maria Pierantoni, Malin Hammerman, Linnea Andersson, Isabella Silva Barreto, Vladimir Novak, Hanna Isaksson, Pernilla Eliasson

## Abstract

Heterotopic ossification (HO) entails pathological mineral formation inside soft tissues. In human Achilles tendons, HO is often associated with tendinopathies, tendon weakness and pain. One hypothesis is that HO occurs in response to inflammation and by either intramembranous ossification, endochondral ossification, or a combination of both. However, refined details regarding HO deposition and microstructure are still unknown. In this study, we characterize HO in intact rat Achilles tendons through high-resolution phase contrast enhanced synchrotron X-ray tomographic imaging. Furthermore, we test the potential of using a procedure to induce local tissue injury by needling to study the relation between microdamage and formation of HO. The results show that HO occurs in all intact rat tendons occupying up to 1% of the total volume at 16 weeks of age. The HOs are characterized by an elongated ellipsoidal shape and by a distinctive fiber-like internal structure which suggests that some collagen fibers have become mineralized. The data indicates that the deposition along the fibers initiates in the pericellular area, and propagates into the intercellular area. The results also show that multiple HO deposits may merge into bigger structures with time by accession along unmineralized fibers. Furthermore, the presence of unmineralized regions within the deposits may indicate that HOs are not only growing, but mineral resorption can also occur. Additionally, phase contrast enhanced synchrotron X-ray tomography allowed to distinguish microdamage at the fiber level due to needling and it could in the future enable to elucidate the relation between local inflammation, microdamage, and HO deposition.

## 1. Introduction

Tendinopathies are common musculoskeletal pathologies [1,2]. Heterotopic mineralization is one possible feature associated with tendinopathies that in some cases can give rise to pain and tendon weakness [3,4]. Heterotopic mineralization can be the result of calcification or ossification: where calcification is characterized by the deposition of calcium salts, and ossification occurs when bone-like deposits form [5]. Calcification and ossification cannot be easily distinguished by medical radiography and the two types of mineralization may occur simultaneously [6].

Heterotopic ossification (HO) is a complex pathologic process commonly documented in tendons [4,7,8]. In humans Achilles tendons, HO is more often observed after a severe trauma, but this could be due to lack of systematic screening in the absence of trauma. Recently, the presence of HO was reported after surgically repaired Achilles tendon ruptures in almost 20% of the patients [9]. We still do not fully understand the underlying mechanisms of HO in tendons, and there are no specific treatments developed to limit HO deposition, aside from surgical removal. The specific biological mechanisms guiding deposition of HO need to be further investigated [10]. Recent studies indicate that inflammation could alter the microenvironment and contribute to the formation of HO [11] and that the accumulation of inflammation-associated cells, e.g. macrophages, can promote HO deposition [12]. HO is currently considered to result from two processes [13]: intramembranous ossification (IMO) and endochondral ossification (EO). IMO occurs when mesenchymal cells condense and differentiate into osteoblasts, whereas during EO mesenchymal stem cells differentiate into chondrocytes forming cartilage which serves as a template and guides the mineralisation process. In Achilles tendons, both EO and IMO have been observed [14]. However, the details regarding HO etiology, morphology and deposition process are still unknown.

Animal models are fundamental to close some of the gaps in our knowledge on the formation of HO-deposits in tendons, specifically rats and mice [7,15–18]. Several animal studies have indicated that HO deposition during tendon healing is dictated by EO [7,19–23]. However, very little is known about HO in intact tendons and the mechanisms guiding the formation of HO-deposits in the absence of trauma remain largely unknown. In this study we use a rat model in combination with high-resolution phase contrast enhanced synchrotron X-ray tomography to increase our knowledge of the process of HO deposition in intact Achilles tendons. The results provide a better understanding on the microstructure and deposition mechanism of HO. Furthermore, since inflammation could be contributing to HO deposition [11,24] we investigate the feasibility of using a needle injury protocol in rat Achilles tendons in combination with high-resolution phase contrast enhanced synchrotron X-ray tomography to study the relation between microdamage and HO formation.

## 2. Methods

### 2.1 Animal model

Female Sprague-Dawley rats, 12 weeks old, (n=16) were used (Janvier, Le Genest-Saint-Isle, France). The rats were kept two per cage, with a light-dark cycle of 12 hours and controlled (55%) and temperature (22 °C). Food was provided *ad libitum*. The rats were used for two separate sets of experiments: 1) characterization of HO-deposits in intact tendons (n=7) where no intervention was performed the Achilles tendon (experiment 1 in table 1), and 2) a needle injury protocol (n=9) to investigate the relation between microdamage and formation of HO-deposits (experiment 2 in table 1). A similar needle injury model was previously performed on rat healing tendons [25,26] and it was used in mice to study HO [4,24]. For the needle injury experiment the rats were divided in 3 groups: a control group (0 needling, n=3), needled 5 times (n=3) and needled 20 times (n=3). Animals were anesthetised with isoflurane. Needle injury was induced by percutaneous penetrations into the mid portion of the right Achilles tendon using an insulin needle (0.25 mm in diameter, size 31G), followed by 5 needle movements inside the tendon tissue in different directions. For the 5 times group, needling was performed through the skin from the lateral side, while the 20 times group was performed through the lateral, medial, proximal, and distal side. This resulted in a total of 20 punctures into the tendon. After four weeks the rats were euthanized and the tendons along with the muscle complex and the calcaneal bone were dissected, placed in a phosphate buffered saline solution (PBS), and frozen at −20° C.

**Table 1.** Quantification of the HO-deposit volumes, volume fractions, and HO locations. All data is presented as mean ± SD. For determining HO-deposit location, the tendon length was normalized so that 0 is the bone junction and 1 is the muscle junction.

| Experiment | Samples | Tendon length (mm) | Number of deposits (tendons) | Volume of individual deposits ($\text{mm}^3$) | Deposit total volume ($\text{mm}^3$) | Volume fraction (%) | HO location (mm) |
| --- | --- | --- | --- | --- | --- | --- | --- |
| 1 | Intact tendons (n=7) | $1.20 \pm 0.08$ | 1(3), 2(2), 3(1), 6(1) | $0.03 \pm 0.02$ | $0.08 \pm 0.04$ | $0.9 \pm 0.6$ | $0.29 \pm 0.13$ |
| 2 | Control (n=3) | $1.21 \pm 0.17$ | 1(2), 2(1) | $0.03 \pm 0.03$ | $0.04 \pm 0.03$ | $0.3 \pm 0.3$ | $0.31 \pm 0.07$ |
| | Needling 5 times (n=3) | $1.21 \pm 0.13$ | 1 (2), 3(1) | $0.07 \pm 0.08$ | $0.10 \pm 0.09$ | $0.7 \pm 0.8$ | $0.25 \pm 0.09$ |
| | Needling 20 times (n=3) | $1.14 \pm 0.11$ | 1 (2), 2(1) | $0.06 \pm 0.04$ | $0.10 \pm 0.01$ | $0.7 \pm 0.2$ | $0.30 \pm 0.10$ |

### 2.2 Phase contrast enhanced synchrotron X-ray tomography (SR-PhC-μCT)

The tendons were mounted in 2 ml Eppendorf tubes filled with PBS, following previous studies [27]. The imaging was performed at the X02DA TOMCAT beamline at the Swiss Light Source (SLS), Paul Scherrer Institute (Villigen, Switzerland) [28]. The images were acquired using a High Numerical Aperture Microscope setup (4x magnification, field of view of 4.2 mm x 3.5 mm and final pixel size of 1.63 µm). The optical setup was coupled to a LuAG:Ce scintillator screen of 150 μm. A propagation distance of 150 mm was chosen, the X-ray energy was set at 15 keV, 2001 projections were acquired over 180° of continuous rotation, and the exposure time was 33ms. The samples were imaged in two-three consecutive locations from the bone junction moving up to muscle junction the tendon. Projections were collected by a pco.edge 5.5 Camera and corrected with dark and flat-field images. The projected density of the sample was calculated using the Paganin phase retrieval for homogeneous objects [29]. The tomographic reconstruction was performed using a Fourier based regridding algorithm [30].

### 2.3 Image processing

To visualize the whole tendons, 2-3 volumes were stitched together using the BigStitcher software package for ImageJ [31]. Then, volume renderings were performed in Dragonfly (v 4.1, ORS software) [32]. The quantitative image analysis was performed in MATLAB. HO-deposits and tendon tissue were segmented by binarization. The threshold values were adjusted for each type of tissue and morphological operations of opening, closing and gap filling were performed iteratively to optimize the segmentation output. The volumes of HO-deposit and total tendon volumes were then calculated by summing the white pixels in each binary segmented image. The volume fraction of HO-deposits defined as the percentage of tendon tissue occupied by HO-deposits was calculated as total volume of HO-deposits divided by the total tendon volume. The location of the deposits was calculated as the distance from the HO-deposit center to the calcaneal bone. To compare the HO-deposit locations between different tendons the distance was normalized to the total tendon length so that the end of the calcaneal bone is equal to 0 and the start of the muscle complex to 1.

## 3. Results

### 3.1 HO in intact tendons

HO-deposits were present in all intact tendons, on the anterior side of the distal third of the tendon (table 1and Fig. 1A). The tendons contained between 1 to 6 individual HO-deposits with an average size of 0.03 ± 0.02 mm^3^, occupying about 1% of the tendon total volume (table 1).

**Figure 1.**
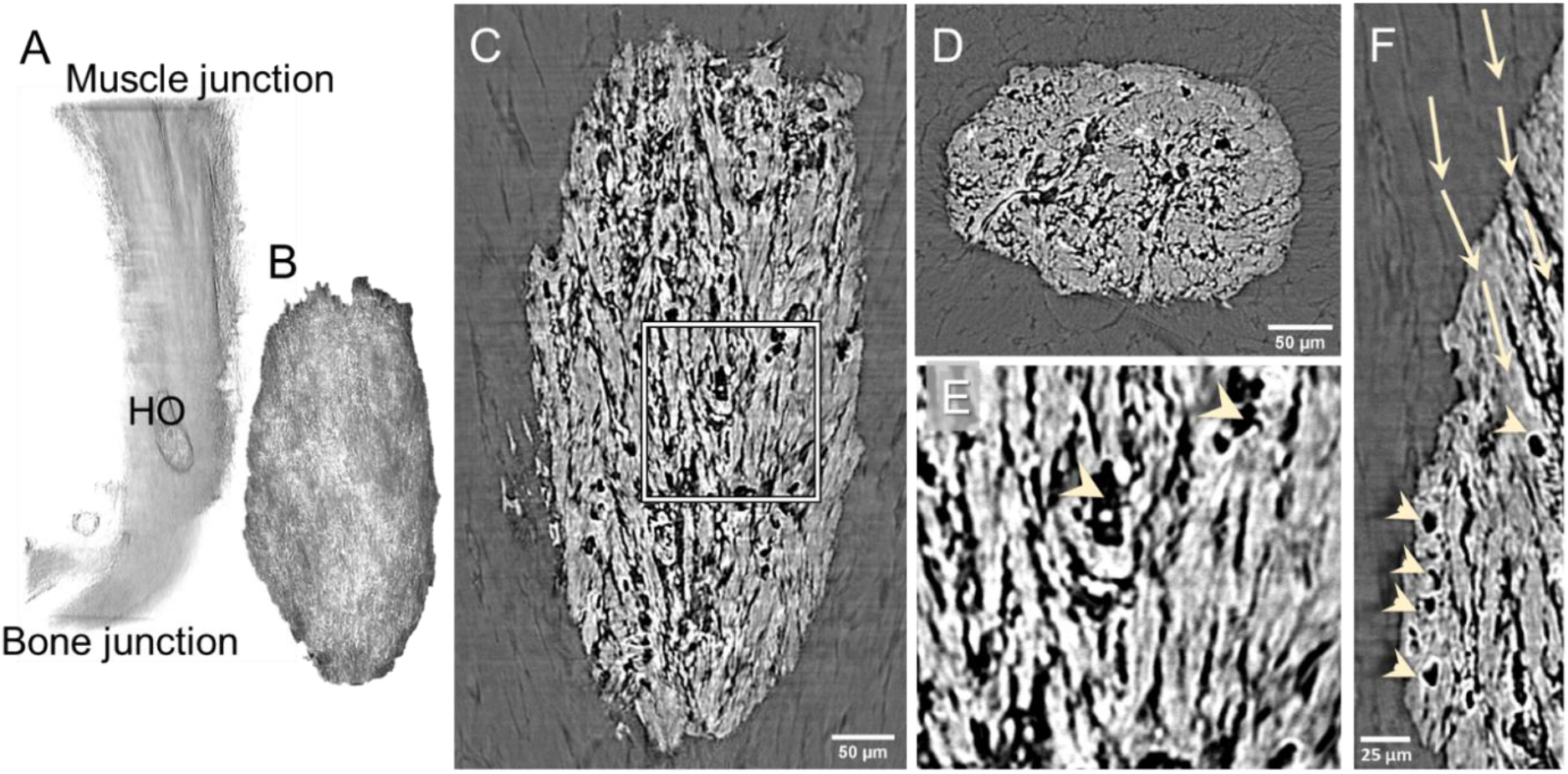
HO-deposits in intact Achilles tendons are characterized by a fiber like-structure. A) volume rendering showing the location of the HO in the whole tendon (from bone to muscle junction), B) volume rendering of one deposit of HO. Longitudinal (C) and cross-section (D) slices showing the internal fiber-like structure of the HO-deposit. E) magnification of the area in the square in C) and F) magnification showing the transition from the collagen fiber to the mineralized fiber in the tendon tissue. Arrowheads indicate cell-like lacunae within the deposits and the arrows show how the fiber structure is maintained moving from the collagen tissue into the mineralized deposit.

All HO-deposits were characterized by an elongated ellipsoidal morphology and a fiber-like internal structure (Fig. 1C-F, video 1). Some of the tendon collagen fibers had mineralized while preserving their structure and could be tracked from the tendon soft tissue into the HO-deposits (Fig. 1F arrows). Within the HO deposits, round unmineralized lacunae of about 15µm diameter could be observed possibly indicating the presence of cells (Fig. 1E, F arrowheads).

Furthermore, in 11 out of 29 (∼40%) of the analyzed HO-deposits large unmineralized gaps were observed within the calcified structure (Fig. 2). In most cases the deposits were located within the tendon margins (Fig. 2A) and unmineralized fibers were visible in the gaps (Fig. 2B). Additionally, adipocyte-like structures were found in some gaps, but only when the minerals bulged out from the tendons (Fig. 2C-D).

**Figure 2.**
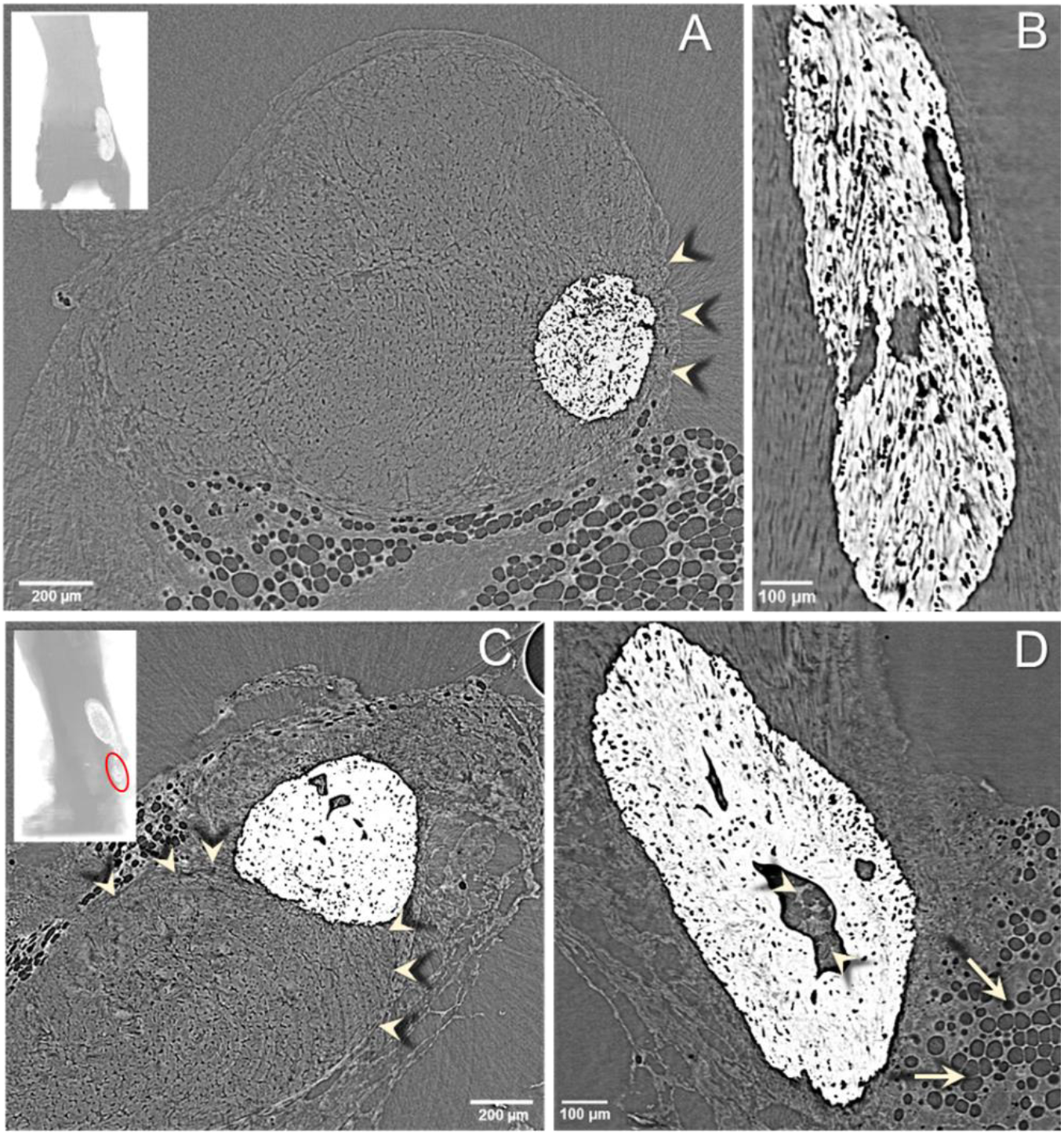
Some HO-deposits contain unmineralized gaps. A) tendon cross section showing the location of the HO-deposit within the tendon margins (arrowheads), B) longitudinal view showing three gaps within one HO-deposit. C) Tendon cross section showing that the HO-deposit bulges out from the tendon margins (arrowheads) into the surrounding tissue. D) Gap within a deposit containing adipocyte-like structures (arrowheads), similar to adipose tissue found outside the tendon (arrows). Inserts: longitudinal views showing the position of the deposits from the calcaneus bone on the bottom (A) anterior, C) lateral views, the circle indicates the deposit shown in the main panel).

### 3.2 HO deposits in micro-injured tendons

Regions where the needle had penetrated the tissue could be identified as microdamage in the collagen fiber structure and HO-deposits were found in the vicinity of the microdamage (Fig. 3).

**Figure 3.**
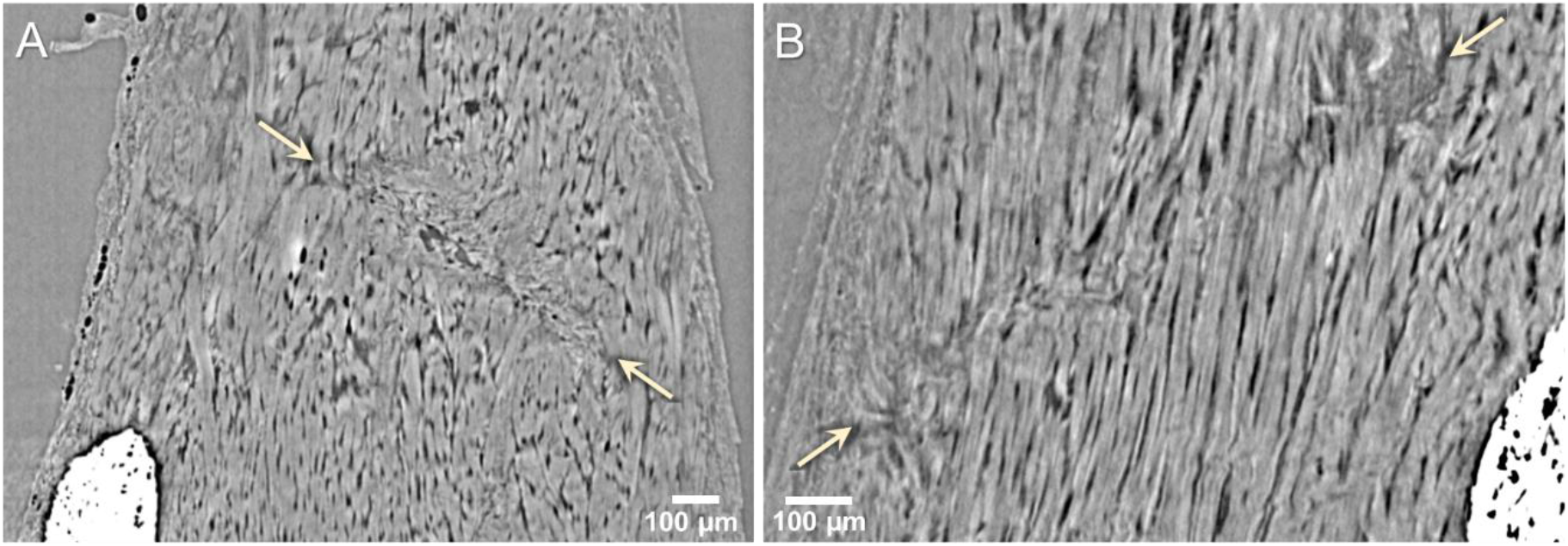
Microdamage in the collagen structure induced by the needling, still visible after 4 weeks. A) 5 times needled sample and B) 20 times needled sample. The arrows indicate the beginning and end of the damaged region. In both tendons HO-deposits in the proximity of the needled regions were observed.

HO-deposits were present in all studied tendons and were located within the distal third of the tendon with one exception where the deposits extended up to almost half of the tendon total length (Fig. 4 and table 1). As in the uninjured tendons, all HO-deposits were characterized by a fiber-like internal structure. The sizes of the deposits were comparable for the two needled groups and possibly slightly bigger than in the control group, occupying about 0.7% of the total volume in the case of the needled tendons and about 0.3% in the case of the controls (table 1). However, the small group size in the needling experiment prevented further statistical analysis. One interesting aspect observed in two of the needled tendons was that multiple HO-deposits were merging. In areas between individual deposits the mineralization seems to proceed along some of the fibers creating points of contact (Fig.4 B-D).

**Figure 4.**
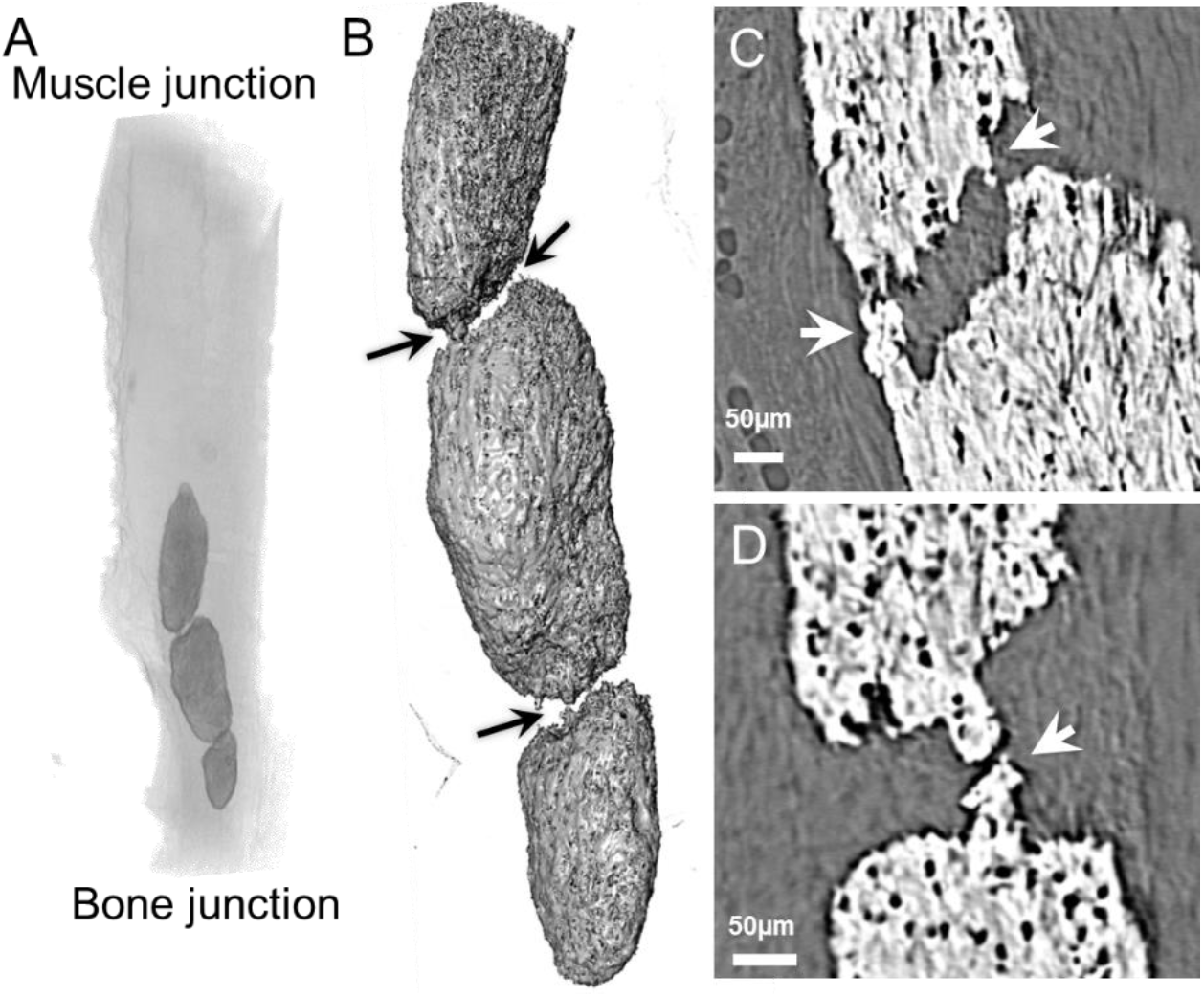
Three merging deposits of HO. A) volume rendering showing the location and B) volume rendering of the HO-deposits. The arrows indicate the merging points. C-D) Longitudinal magnifications showing the merging between the top and middle deposits (C) and between the middle and bottom deposit (D).

Moreover, in one of the needled samples early stages of HO could be identified (Fig. 5, video 2). The HO occurred at about ½ of the tendon lengths at the lateral edge of the tendon, close to where the needle was inserted, however microdamage from the needling was not visible in this sample. The data seem to indicate that mineral deposition starts around cells (arrowhead) and propagates along fibrils and fibers in between them (arrow).

**Figure 5.**
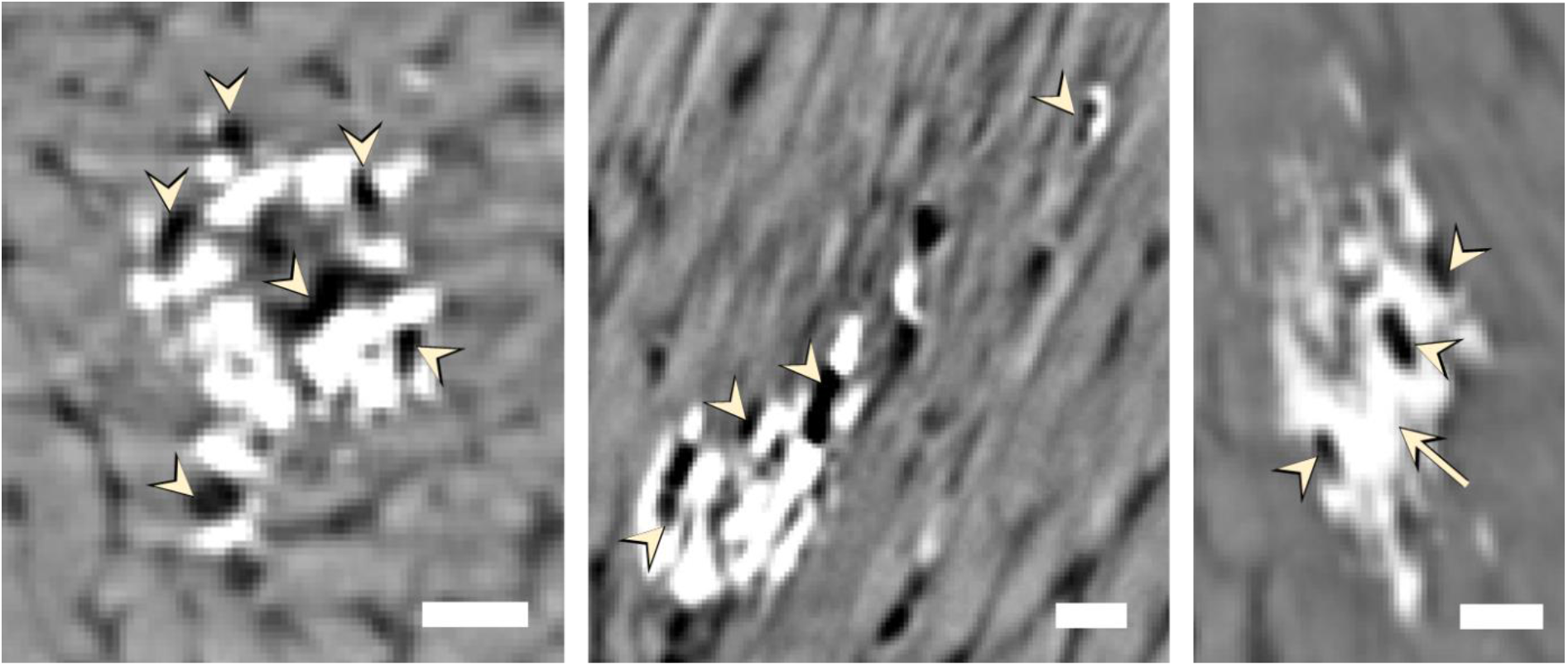
Forming HO-deposits in a 5 times needled sample. From left to right: cross-sectional, left and right views of the deposit showing that HO may start around tenocyte cells (arrowheads) and propagates along the fibers in between them (arrowhead). Scale bars = 25 µm.

## 4. Discussion

In this study the microstructure of HO-deposits in intact rat Achilles tendons was characterized in detail. Deposits of HO were present in all tendons, had an elongated ellipsoidal shape, and occupied up to 1% of the tendon volume. They were mostly located on the anterior distal part close to the tendon-surface, in some cases protruding out from the tendon. The HO deposits were characterized by a fiber-like structure in which some fibers from the tendon soft collagen matrix had become mineralized. The structure of the deposits strongly resembles the one of calcified avian tendons where the calcification occur hierarchically along the collagen fibrils, fibers and bundles [33,34]. Questions of how collagen fibers are mineralized into hard tissue in intact tendons and what factors trigger this process remain open. However, the fiber-like structure indicates that it primarily is through IMO. Several previous studies have shown that HO in healing tendons primarily occurs through EO [7,20–23]. These results for healing tendons and our findings in intact tendons together indicate that HO could possibly occur through different pathways in intact and injured tendons.

In vitro studies have shown that tenocytes have a predisposition to differentiate into chondrocyte-like cells that produce calcium deposits [35]. In our study small ellipsoidal gaps were distinguished within the OH-deposit structure. These gaps (or lacunae) may indicate the presence of inflammatory cells such as macrophages or masts that promote HO deposition [12]. However, the lacunae could also consist of chondrocytes or tenocytes, as suggested by their locations, sizes, and organization [27,36]. A recent study has shown that in naturally mineralizing turkey tendons, calcification occurs along fibrils in the correspondence of canaliculi which are radiating from tenocytes [37]. The results here reported for early stages of HO (Fig. 5) could indicate that also in intact rat tendons mineral deposition could be associated with tenocytes. Further investigation in this direction could provide fundamental insight into HO. Additional studies may improve our understanding of pathological ossification mechanisms in vertebrate tissues by expanding the notion of the existence of a close association between the calcification process, tenocytes/lacunae, and collagen fibrils/fibers from philological mineralization in avian tendons to pathological deposition in mammal intact tendons.

Further information about the deposition process was provided by the observation of large unmineralized areas in about 40% of the HO-deposits. The presence of these unmineralized areas could either be due to how the mineral grew during the deposition process or it could also indicate that mineral reabsorption occurs inside the deposits. In the case of HO-deposits bulging out of the tendon, some of the unmineralized areas spherical structures reassembling the adipose tissue on the outside of the tendon were visible (Fig. 2D). This could indicate that during mineral deposition some of the adipose tissue has been encapsulated within the HO deposits.

This study also investigated the feasibility of using needling of the rat tendon to study the relationship between microtrauma and HO. The data may indicate that the volume fraction of HO-deposits in the needled groups appears to be higher than in the control group. Specifically, since the deposited HO-deposits were close to the microdamage. One further interesting observation for needled tendons was the merging of multiple HO-deposits by mineral accretion along some of the non-mineralized fibers in between deposits. This could indicate that with time one single bigger structure may be formed from the merging deposits which could affect the mechanical performance of the tendon substantially [24].

Species specific differences exist in HO deposition in tendons [4]. For instance, mice develop tendon mineralization with ageing while rats develop HO from a young age. Consequently, in previous studies, mice in the control group did not show any big HO-deposits, whereas in the case of this study all the intact rat tendons contained HO-deposits. The presence of pre-existing deposits can make it harder to identify the effect of the needling in rats. In future, further investigation using younger rats could help understanding when HO deposition starts in intact rat Achilles tendons. Furthermore, using younger rats, and consequently having a control group totally lacking HOs, could possibly provide a clearer understanding of the relation between microdamage and HO formation.

Furthermore, in the attempt to capture the very early stages of HO deposition, the rats were sacrificed only 4 weeks after the needling, whereas in previous studies HO formation was reported in needled mouse tendons after 20 weeks [4,24]. Considering that new HO deposition could be identified in only one of the samples, 4 weeks may not be sufficient for the mineral deposition to initiate. Consequently, it could be beneficial to extend the time between the needling and the rat sacrifice to assess a definitive correlation between the needling and new HO deposition.

Regardless of the differences discussed above, similarities between the mouse and rat models can be noted. Both in mice and rats HO-deposits had a similar overall shape and were identified only close to the calcaneus bone. Furthermore, in both models some of the HOs were observed very close to the tendon surface, almost bulging out from it.

## Conclusions

Heterotopic Ossification is associated with tendinopathy and may result in discomfort and impaired mechanical properties. Our results show that HO deposits are present in intact rat tendons, and that they are characterized by a unique structure in which some fibers from the tendon soft collagen matrix have mineralized. Moreover, the HO deposition seems to initiate around cells, possibly tenocytes, and propagate along the fibers in between them. Furthermore, we show that using a needle injury animal model in combination with phase contrast enhanced synchrotron X-ray tomography is a promising approach to determining the relation between local microdamage at the collagen fiber level and HO formation.

## Acknowledgments

Funding from the Knut and Alice Wallenberg KAW Foundation (Wallenberg Academy Fellows 2017.0221) and the European Research Council (ERC) under the European Union’s Horizon 2020 research and innovation programme (grant agreement No 101002516), the Swedish research council (2017-00990), the Royal Physiographic Society of Lund and the Greta and Johan Kocks foundation, the Swedish National Centre for Research in Sports (P2021-0032)), Magnus Bergvall foundation (2019-03211) and Åke Wiberg foundation (M20-0021) are greatly acknowledged. VN acknowledges funding from the European Union’s Horizon 2020 research and innovation program under the Marie Skłodowska-Curie Grant Agreement No 701647. We acknowledge the Paul Scherrer Institut, Villigen, Switzerland for provision of synchrotron radiation beamtime at the TOMCAT beamline X02DA and of the SLS.

